# Recreating the Native Airway Microenvironment Using Tissue-Specific Extracellular Matrix Bioinks for Proximal Airway Engineering

**DOI:** 10.64898/2026.06.04.730194

**Authors:** Heather Wanczyk, Nina D. Kosciuszek, Joanne Walker, Daniel J Weiss, Christine Finck

## Abstract

*Ex vivo* airway engineering approaches such as 3D bioprinting offer a promising strategy for generating functional airway replacements, but the fabrication of hollow, patient-specific proximal airway constructs using translationally relevant bioinks remains challenging. This study describes the development of biocompatible, polymer-blended human airway-derived decellularized extracellular matrix (AW-dECM) bioinks for engineering structurally and mechanically relevant airway tissues. An optimal formulation consisting of 30 mg/mL AW-dECM and nanofibrillar cellulose alginate conjugated to RGD supported the bioprinting of simple and complex hollow airway structures with mechanical properties comparable to native airways (∼8–10 kPa). The bioinks also promoted primary human airway epithelial cell viability, adhesion, and differentiation into mucociliary and secretory phenotypes during 28 days of air-liquid interface culture. Furthermore, subcutaneous implantation in immunocompetent rats demonstrated excellent biodegradative stability and overall biocompatibility over 30 days. Collectively, these findings establish a foundation for improved physiological airway models and future tissue-engineered airway replacements.

## Introduction

*Ex vivo* bioengineering approaches for proximal airways remain challenging despite decades of research ^1–4^. Emerging technologies like 3D bioprinting are now being applied to generate tracheal and airway constructs for congenital pediatric respiratory defects and for acute or chronic airway diseases or injury ^5–8^. These scaffolds, combined with stem cells such as primary bronchial epithelial cells or human induced pluripotent stem cells differentiated to lung progenitors, have the potential to regenerate airway epithelium and restore tissue structure following injury. As current treatments for *congenital airway defects* (i.e., laryngeal clefts ^9^, tracheomalacia, bronchomalacia ^10^) or *respiratory diseases* (i.e., cystic fibrosis (CF) ^11^ and primary ciliary dyskinesia (PCD) ^12^, fail to regenerate damaged or defective airway tissue, this approach would significantly improve available therapeutic options.

An optimal approach would pair stem cells with biologically relevant scaffolds capable of remodeling and growing *in vivo*. Previously used cell types for proximal airway modeling and engineering applications include human primary bronchial epithelial cells (hBECs) or induced pluripotent stem cells (iPSCs) differentiated to basal airway epithelial progenitors that, when grown at air-liquid-interface, differentiate into a mature pseudostratified epithelium complete with functional mucociliary and secretory cells ^13–16^. Substrates commonly used in these approaches include synthetic porous membranes such as polyethylene terephthalate (PET) coated with purified extracellular matrix (ECM) proteins like Collagen or Matrigel^TM^ ^17^, which fail to recapitulate the stiffness, viscoelasticity and biochemical complexity of native airways. This can lead to disparate cell morphologies and transcriptional profiles, significantly limiting the clinical relevance of findings. 3D bioprinting can overcome these limitations by enabling the precise control and fabrication of complex, physiologically relevant constructs in a layer-by-layer fashion that better mimic native airway mechanics and architecture. Among candidate materials, ECM is particularly promising due to its complex composition of proteins, glycoproteins and growth factors that regulate cell behavior and tissue homeostasis. Unlike synthetic materials that are biologically inert, ECM-based materials have the potential to better support airway progenitor cells and enable ongoing physiological remodeling, making them especially suitable for pediatric applications.

In several studies, decellularized ECM-based (dECM) materials have been demonstrated to enhance the differentiation and function of seeded or embedded iPSC-derived lung progenitors or primary airway/alveolar epithelial cells ^18–21^. To render the tissue immunologically inert, these materials are processed using detergents such as Sodium Dodecyl Sulfate (SDS) or Sodium Deoxycholate (SDC) to remove all cellular and antigenic components ^22^. dECM is preferred over synthetic sources such as poly (lactic-co-glycolic) acid (PLGA) and polyethylene glycol (PEG) as it retains essential biochemical constituents, ECM proteins and mechanical properties most akin to native tissue that help to promote appropriate cell adhesion, growth and differentiation. Additionally, dECM-based materials have the potential to integrate into host tissue and remodel appropriately over time ^23^, which is particularly important for the development of tissue engineered constructs for pediatric patients that are still growing ^24–27^.

Recent studies using dECM-based bioinks for reconstructing kidney ^28, 29^, liver ^30–32^, GI ^33^ and heart ^34–36^ tissues have shown promising results in pre-clinical models. However, our recent work (unpublished) has demonstrated that airway dECM has a low stiffness (∼100-300 Pa), which makes it difficult for these soft bioprinted structures to maintain their shape following gravity-induced deformation. Therefore, this material is typically combined with other bio-compatible polymers such alginate, gelatin or synthetic materials (i.e., polycaprolactone or PLGA) to increase its mechanical properties. Subsequently, this also increases the shear thinning behavior of blended dECM materials, which is a property required for 3D bioprinting different materials with high fidelity ^37^. To allow for hollow organ fabrication, yield-stress support baths have also been developed where bioinks can be directly extruded to support complex tissue design. One novel approach called freeform reversible embedding of suspended hydrogel (FRESH), involves the use of a gelatin microparticle slurry bath whereby bionks can be directly extruded into support baths containing ionic molecules to facilitate crosslinking, gelation and development of complex, hollow 3D scaffolds ^38^.

The following work demonstrates the development and optimization of regionally isolated human airway derived decellularized ECM blended with alginate, nanofibrillar cellulose-RGD (Cellink-RGD) conjugated bioinks for 3D bioprinting hollow airway structures. Rheological assessments demonstrate that AW-dECM + Cellink-RGD bioinks exhibit mechanical properties within the range of native airway tissue (Young’s modulus=∼6-8 kPa) as well as excellent printability and shape fidelity. In combination with the FRESH approach, we demonstrate feasibility for bioprinting both simplified and patient-specific hollow airway structures with high resolution and reproducibility. Additionally, primary airway epithelial cells seeded on top of bioinks and cultured at ALI exhibit excellent viability and differentiation to mucociliary and secretory phenotypes over 28 days. Finally, subcutaneous implantation of AW-dECM + Cellink-RGD bioinks in rats reveal excellent stability and biocompatibility over 30 days. Overall, this study provides proof of concept for the development of human AW-dECM blended polymer bioinks for 3D bioprinting complex airway structures with excellent mechanical and biochemical properties, while also further informing on host immune response to commercially available alginate-RGD based biomaterials. This information will be pertinent in the improved design of translationally relevant materials useful for airway modeling and regeneration.

## MATERIALS AND METHODS

### 1.1. Human lung decellularization and airway-specific isolation

Human lungs from patients (2 donors) with no known lung diseases or smoking history were obtained from the Autopsy Services at the University of Vermont (UVM) and were decellularized using established protocols ^39, 40^. Adequacy of decellularization was demonstrated by lack of intact cells or cell debris on hematoxylin and eosin-stained mounted 5 µm paraformaldehyde-fixed sections, no intact bands on DNA gels, and < 50 ng DNA/ug protein ^39, 40^. Airway-specific regions were then isolated from decellularized lungs using a previous protocol [26]. To produce the hydrogel, airway preparations were cryomilled, lyophilized and digested under constant mixing over 72 h using 1 mg/mL pepsin (ThermoFisher Scientific) + 0.01 M HCl (Fisher) ^40, 41^. Insoluble aggregates were removed by centrifuging the mixture at 600 rpm for 20 minutes, followed by removal of the supernatant. Digested lung tissue was pH balanced to 7.0 by adding 1:9 the volume of 0.1M NaOH to the mixture. Samples were then frozen at -80℃ for 24 hours and lyophilized for an additional 72 hours. The decellularized lung tissues and subsequent hydrogel preparations have previously been extensively characterized for protein and glycosaminoglycan contents ^39^.

### 1.2. Preparation of AW-dECM polymer-blended bioinks for mechanical studies

Polymer-blended AW-dECM bioinks were prepared by solubilizing 45 mg lyophilized airway tissue in 1.5 mL Cellink-RGD bioinks (Cellink). Briefly, lyophilized lung tissue was added to a 3 mL syringe and mixed with Cellink-RGD. The bioink suspension was mixed using two 3 mL syringes connected via a female-female luer lock until a homogenous solution was obtained. Solubilized bioinks were then placed on ice for at least 1 hour. For comparison, 45 mg lyophilized AW-dECM was also resuspended in 1.5 mL cold Dulbecco’s Modified Eagle’s Medium (DMEM) (Fisher) and placed on ice for solubilization over 1 hour.

### 1.3. Oscillatory rheology measurements of AW-dECM Cellink-RGD bioinks

To assess the mechanical properties of 30 mg/mL AW-dECM alone and 30 mg/mL AW-dECM combined with Cellink-RGD bioink, a DHR-3 Discovery Rheometer (TA Instruments) was utilized. 600 uL of each of crosslinked, previously swollen in PBS, hydrogel sample (n=6 per hydrogel type) was loaded onto a 12 mm parallel steel plate pre-warmed to 37°C. Gap height was set to 1000 µm with 1% strain and 0.159 Hz frequency (1.0 rad/s). Oscillatory frequency sweeps were performed at 37°C on all materials and the storage modulus (G’) and loss modulus (G’’) were determined. To assess the shear thinning behavior (printability) or viscosity of AW-dECM + Cellink-RGD bioinks, flow sweep tests were performed on uncrosslinked samples (n=6) using a 12 mm parallel steel plate set to 25°C, gap height= 1200 µm and shear rate = 0.001s to 200s.

### 1.4. SEM analysis of bioink preparations

30 mg/mL AW-dECM alone or blended with Cellink-RGD bioink were prepared for scanning electron microscopy analysis. Briefly, 3D bioprinted and crosslinked samples (8 mm sized discs, 2 mm thickness) were fixed with 2.5% Glutaraldehyde in 0.1 M cacodylate buffer at room temperature for 1 hour. For post fixation steps, samples were submerged in 1% OxO4, 0.8% Ferricyanide in 0.1 M cacodylate buffer at room temperature for 1 hour. Samples were then rinsed 3x with 0.1 M cacodylate buffer, followed by rinsing with distilled water (3x). Samples were dehydrated with ethanol for over 40 minutes and dried using 1:2 solution of hexamethyldisilazane (HMDS) and 100% ethanol, followed by a 2:1 solution of HMDS and ethanol for 20 minutes each. For final steps, samples were rinsed in 100% HDMS for 20 minutes (x2). Samples were then sputter coated with gold and imaged using a JEOL JSM-5900LV scanning electron microscope.

### 1.5. Fabrication of 3D bioprinter from commercially available FDM printer

A Flash Forge Adventurer 5M desktop printer (Flashforge) was modified into a 3D bioprinter based on a concept developed by researchers at Carnegie Mellon University and demonstrated at the “3D Bioprinting Open-Source Workshop” ^42^ (**Supplementary Figure 1**). Briefly, Computer-Aided Design (CAD) files were uploaded and 3D printed to assemble the syringe pump (Replistruder) required for layer-by-layer deposition of bioinks (**Supplementary Figure 1A**). The original parts were removed from the printer, and the 3D printed Replistruder parts were assembled (**Supplementary Figures 1B + 1C**). Next, the original wifi board was removed and replaced with a Duet Wifi board following a previous protocol ^42^ (**Supplementary Figure 1D**).

### 1.6. Conversion of pediatric airway CT scan into 3D printable file

Using an open-source online medical data repository (embodi3D), a CT scan of a pediatric airway was acquired and corresponding DICOM files were uploaded into the imaging platform Slicer 4.11. No HIPAA-protected information was obtained or utilized for the scan. Files were converted into .stl files and modified in Solidworks. The converted airway model was then uploaded into PRUSA software whereby it was modified and converted into G code for subsequent bioprinting (**Supplementary Figure 2**). Final dimensions were 65 mm x 60 mm x 27 mm with 100% rectilinear infill.

### 1.7. FRESH support preparation

FRESH v2.0 gelatin microparticle support bath solutions were prepared using a complex coacervation method based on a previous protocol ^43^. Briefly, 125 mL distilled water was added to a 500 mL glass beaker and heated to 50°C, followed by the addition of 125 mL 100% ethanol (Fisher Scientific). To this solution, 3% Gelatin B (Fisher), 0.3% Gum Arabic, and 0.125% Pluronic were added stepwise. A stir bar was placed into the solution and the beaker covered with parafilm, followed by mixing at 350 rpm overnight. The following day, stirring was stopped to allow settling of the formed gelatin microparticles and after ∼30 minutes, excess ethanol was removed. Gelatin microparticles were collected in a 50 mL conical tube and spun down at 750 xg for 3 minutes. Excess solution was decanted followed by rinsing, mixing samples with distilled water and spinning at 1000 xg for 3 minutes (x2). For the final step, the support bath was mixed with 0.1% CaCl_2_, spun at 1000 xg for 3 minutes and then set aside for subsequent printing. All support bath materials were used within a 2-hour period to prevent material dehydration.

### 1.8. Extrusion-based 3D bioprinting of patients-specific airway and simplified hollow airway structure using AW-dECM polymer-blended bioinks and FRESH support

Simplified hollow airway structures (50 mm x 40 mm x 20 mm) were created using SolidWorks. Additionally, hollow cylinders (12 mm diameter x 30 mm length), one with a wall thickness of 2 mm and the other 1mm, were designed using Tinkercad. 3D models were uploaded into PRUSA software and converted into G-codes reflecting diameter of the syringe and needle used for bioprinting. These files were then uploaded into the custom built bioprinter. All prints were made using a 30G needle (120 µm) with a layer height set to 0.06 mm and 3 perimeters. For support bath setup, FRESH slurry was added to a 60 mm tissue culture dish and set up for bioprinting. Prepared bioinks were put into a 3 mL Hamilton gas tight syringe and loaded into the Replistruder syringe pump for FRESH bioprinting. Following extrusion of inks within the gelatin microparticle support bath, the airway scaffolds were released by incubating the print containers at 37°C for 1 hour.

### 1.9. Culture and seeding of primary human airway epithelial stem cells on 3D bioprinted AW-dECM + Cellink-RGD bioinks

Primary human airway epithelial cells (P1) were graciously provided by the Ryan Lab at the University of Iowa State. 500k cells were cultured and expanded in T75 flasks (Fisher) using Pneumacult Ex Medium (Stemcell Technologies) + supplements (A0830, DH1, Hydrocortisone, Rock Inhibitor). Airway cells were then seeded at 50k/bioprinted discs (6 mm x 1 mm thick, crosslinked for 1.5 hrs in 0.1% CaCl_2_ and rinsed 2x with PBS and incubated at 37°C for 1 hr) in transwell inserts and cultured in Pneumacult-EX medium over 4 days. Cells were then air lifted and differentiation medium (Pneumacult ALI-Stemcell Technologies) + supplements (Hydrocortisone, Heparin) were added to the basal compartment of the wells (n=12 wells/cell line). Cells were then harvested on Day 28 for qPCR analysis.

### 1.10. *In vitro* assessment of airway epithelial cell viability on 3D bioprinted AW-dECM + Cellink-RGD bioinks

To assess biocompatibility of airway cells seeded on 30 mg/mL AW-dECM + Cellink-RGD bioinks, samples were incubated with Calcein AM/Ethidium Bromide staining (Fisher) at 37°C for 10 minutes on Days 1, 3, 7 and 14 (n=3/day). Samples were evaluated using a Zeiss LSM 880 microscope.

### 1.11. qPCR analysis of airway cells seeded on 3D bioprinted AW-dECM + Cellink-RGD bioinks

On Day 28, airway cells seeded on bioinks (n=8) were treated with 1 mL Trizol/well per manufacturer’s protocol. Following cell lysis, 200 uL chloroform was added to each sample, vortexed and incubated at room temperature for 2-3 minutes. Samples were then spun down at 12,000 xg for 15 minutes at 4°C. Following centrifugation, the top phase was removed and 500 uL isopropanol was added and mixed. Samples were incubated at room temperature for 10 minutes and then centrifuged as above for an additional 10 minutes to precipitate isolated RNA. Following centrifugation, the supernatant was removed and 350 uL RLT lysis buffer was added to the RNA pellet. 350 uL 70% ethanol was added to the sample, mixed and dispensed into a column for RNA purification (RNeasy Minikit-Qiagen). RNA concentration was determined using a Nanodrop spectrometer (Thermofisher). Housekeeping genes *GAPDH* and *YWHAZ* were used for normalization of gene expression.

### 1.12. Subcutaneous implantation of acellular 3D bioprinted AW-dECM + Cellink-RGD bioinks in rats

To assess biocompatibility and degradation of bioinks, bioprinted samples (Cellink-RGD alone-**control** or 30 mg/mL AW-dECM + Cellink-RGD-**treatment**; 8 mm x 3 mm discs) were implanted subcutaneously (**contro**l-left side, **treatment**-right side) in the lateral dorsal region of Sprague Dawley rats, weighing 300-500 g (16 weeks of age, n=24; 14 females, 10 males). Prior to surgery, rats were kept in a temperature-controlled environment (22°C) with 50% relative humidity and 12-12-hour light-dark cycles, with access to water and food as needed. Anesthesia was administered using 4% Isoflurane via a nose cone. Two 3-cm incisions were made, one on the left side and one on the right side of the lateral dorsum in each rat under sterile conditions. Each animal received one sample of Cellink-RGD bioink and one sample of 30 mg/mL AW-dECM + Cellink-RGD bioink circular discs. Bioinks were implanted in a subcutaneous pocket between the skin and muscle of each rat and closed with 3-0 Vicryl suture. On Days 3, 14 and 30, eight rats per timepoint were euthanized via CO_2_ overdose to evaluate biocompatibility (fibrosis, inflammation and presence of granulation tissue), angiogenesis and degradation.

### 1.13. Histological Analysis

Tissue samples were fixed in 10% Formalin (Fisher Scientific) for 24 hours. Samples were then embedded in paraffin and 5 µm or 10 µm sections were cut using a microtome. For histological assessments, samples were mounted on positively charged slides (Fisher) and stained with hematoxylin-eosin (H & E) and Masson’s trichrome (MT) stain to evaluate cellular infiltration and fibrosis, respectively. Six fields per scaffold were scanned and quantitative analysis of cellular infiltration (6 images/sample, 10 measurements/image) were performed using ImageJ software. Presence of fibrotic capsules was assessed using MT staining.

### 1.14. Statistical Analysis

Statistical analysis was performed on H & E samples for the evaluation of cell infiltrate density/thickness using One-way Anova with Tukey’s Multiple Comparisons (6 images/sample, 10 measurements/image). A *p*-value<0.05 was considered statistically significant. Data is presented as the mean ± standard error of the mean (SEM). Analyses were performed using GraphPad Prism.

Statistical significance of qPCR data obtained from airway cells seeded on 30 mg/mL AW-dECM compared to 30 mg/mL AW-dECM + alginate-RGD blended bioinks were assessed using One-way Anova with Tukey’s Multiple Comparisons (GraphPad Prism). Housekeeping genes used for normalization were *YWHAZ* and *B2M*. Mean ± SD. p<0.05 *; p<0.01 **; p<0.001 ***; p<0.0001 ****, n=4. Data calculated using delta-delta Ct method. Statistical analysis performed on fold gene expression values (2^-(ΔΔCt).

## 2. RESULTS

### 2.1. AW-dECM polymer-blended RGD bioinks exhibit enhanced mechanical properties compared to AW-dECM alone

The topography and mechanical properties of 30 mg/mL AW-dECM + Cellink-RGD bioink samples were assessed using scanning electron microscopy (SEM) and oscillatory rheometry (**Figure 1**). Analysis of bioink topography using SEM showed that AW-dECM + Cellink-RGD bioinks had a smooth and less porous surface compared to 30 mg/mL AW-dECM alone (**Figure 1A + 1B**). Collagen fibers were evident in both samples (**Figure 1B**). Flow sweeps performed on AW-dECM + Cellink-RGD bioinks demonstrated excellent shear thinning behavior as indicated by a decrease in viscosity with increasing shear rate (∼15 kPa at 0.01 s compared to ∼1.0 Pa at 200 s) (**Figure 1C**). Frequency sweeps also revealed that both materials exhibited viscoelastic properties with increasing strain. 30 mg/mL AW-dECM + Cellink-RGD and Cellink-RGD alone exhibited a statistically significant higher storage modulus (∼8 kPa) compared to 30 mg/mL AW-dECM alone (∼100 Pa) **(Figure 1D + 1E)**.

**Figure 1:**
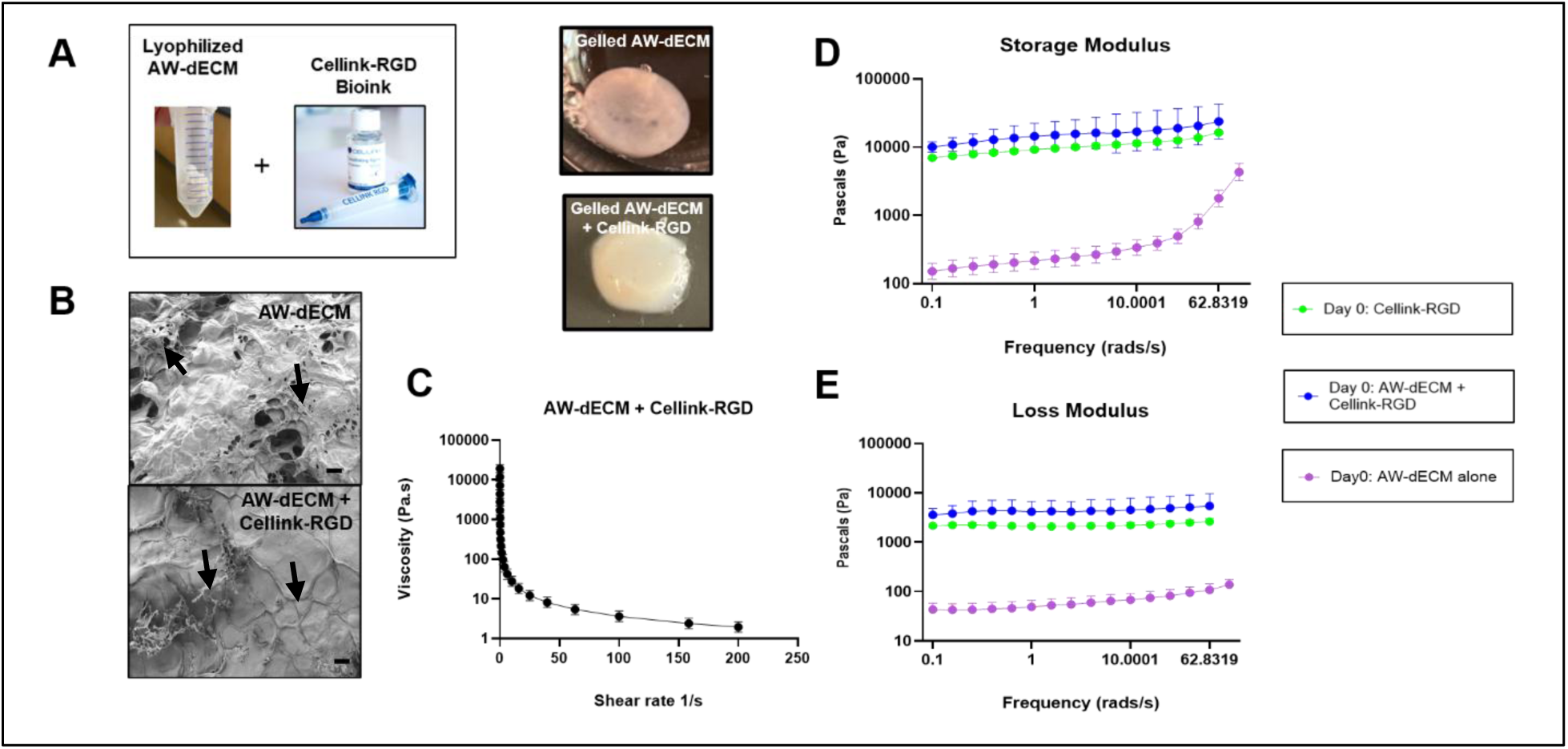
Mechanical and Physical Characterization of AW-dECM ± Cellink-RGD bioinks. **1A:** Macroscopic images of gelled AW-dECM alone and gelled AW-dECM + Cellink-RGD prior to SEM analysis. **1B:** SEM images of AW-dECM vs. AW-dECM + Cellink-RGD, scale bars= 200 µm. Black arrows indicate presence of collagen fibers. **1C**: Flow sweep evaluating viscosity of AW-dECM + Cellink-RGD bioinks in Pa.s, n=6. **1D and 1E**: Frequency sweep evaluating storage and loss modulus in Pascals (Pa) of AW-dECM alone vs. Cellink-RGD alone vs. AW-dECM + Cellink-RGD. Data represented as mean ± SD, n=6 per material type tested.

### 2.2. AW-dECM polymer-blended bioinks can be successfully used to bioprint simplified and patient-specific hollow airway structures

To assess printability and shape fidelity of 30 mg/mL AW-dECM + Cellink-RGD bioinks, hollow cylinders (8 mm diameter x 2 mm height x 2 mm thickness) were bioprinted using the FRESH approach. After preparing the gelatin microparticle slurry bath (**Figure 2A-2D**), extrusion multiplier rate settings (flow rate) ranging from 0.55-1.3 were modified using PrusaSlicer 2.8.0 software prior to printing to determine optimal rate for filament integrity and strength. Five hollow cylinders, each bioprinted using a different flow rate, were extruded into FRESH support baths and removed by placing the printed cylinders at 37 °C for 30 minutes (**Figure 2E**). Based on subsequent print height, rate of bioink shrinkage after 2 hours at room temperature and structural integrity as verified by bright field microscopy, it was determined that a flow rate of 1.3 resulted in the most mechanically stable filament structure (**Figure 2E-2G**). Using these settings, hollow cylinders representing the thickness of a child’s upper airway (20 mm x 5 mm x 1 mm and 20 mm x 10 mm x 2 mm in size), were successfully bioprinted and removed from the bath following incubation at 37°C for 30 minutes. Measurement of the cylinders confirmed that they matched the dimensions of the original CAD files and were able to under manipulation without collapsing (**Figure 2H-2M**). Next, we evaluated the ability to bioprint more complex hollow airway structures using AW-dECM + Cellink-RGD bioinks (**Figure 3A**), which included two simplified hollow airway structures (**Figure 3B + 3C**) and a patient-specific airway derived from a CT scan (**Figure 3D**). The intended hollow geometries were accurately bioprinted with high fidelity and resolution into FRESH support baths using a 30g needle and rectilinear infill pattern set at 100%. Following printing, the airway constructs were successfully removed by incubating the bath at 37° C for 30 minutes to 1 hour.

**Figure 2:**
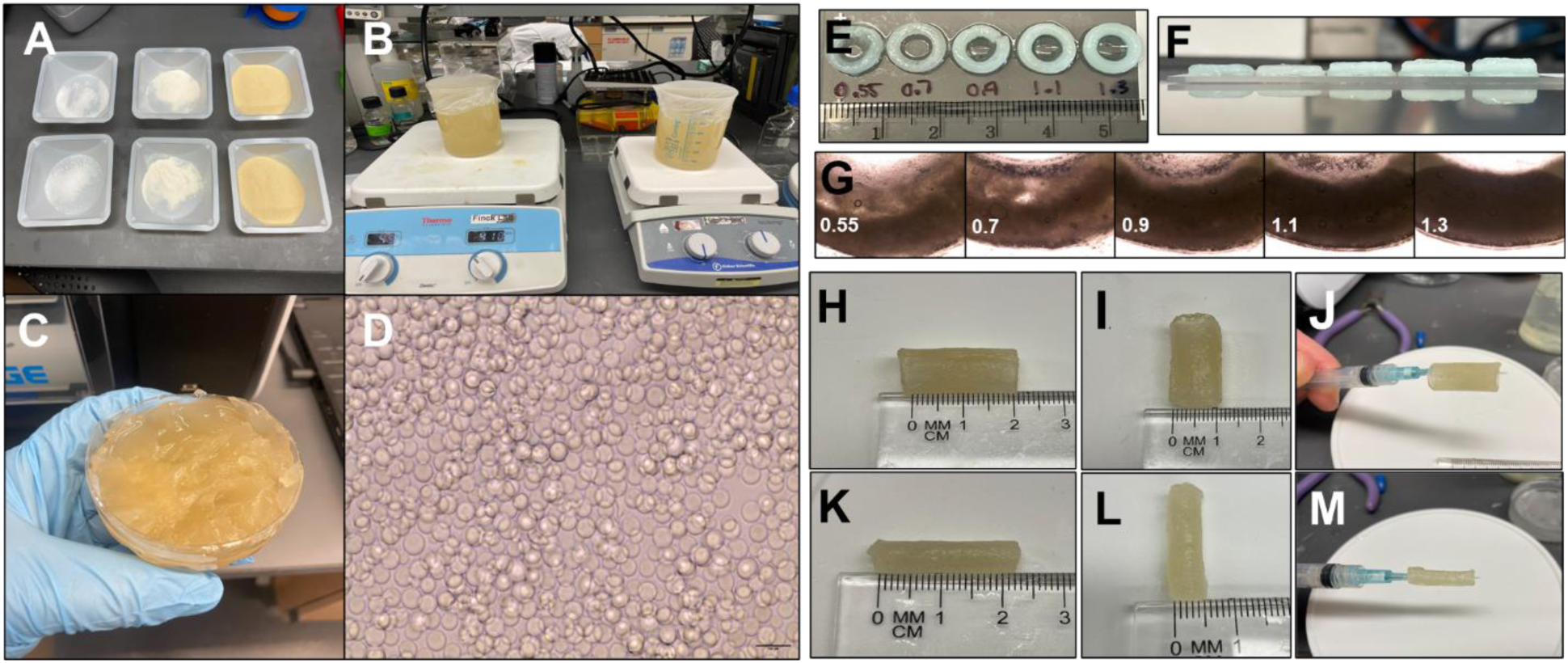
FRESH support preparation and proof of concept for 3D bioprinting hollow cylinders using AW-dECM + Cellink-RGD bioinks. **2A-2D:** Preparation of FRESH support and bright field image of uniform gelatin microparticles. Scale bar = 50 µm. **2E+2F**: Testing different extrusion flow rates for optimal filament density and construct height/thickness. **2G**: Bright field images of hollow, cylinders bioprinted using different extrusion flow rates. **2H-2J:** 20 mm x 10 mm x 2 mm hollow cylinder bioprinted using FRESH approach + 30 mg/mL AW-dECM + Cellink-RGD bioink. **2K-2M**: 20 mm x 5mm x 1 mm hollow cylinder bioprinted using FRESH approach + 30 mg/mL AW-dECM + Cellink-RGD bioink.

**Figure 3:**
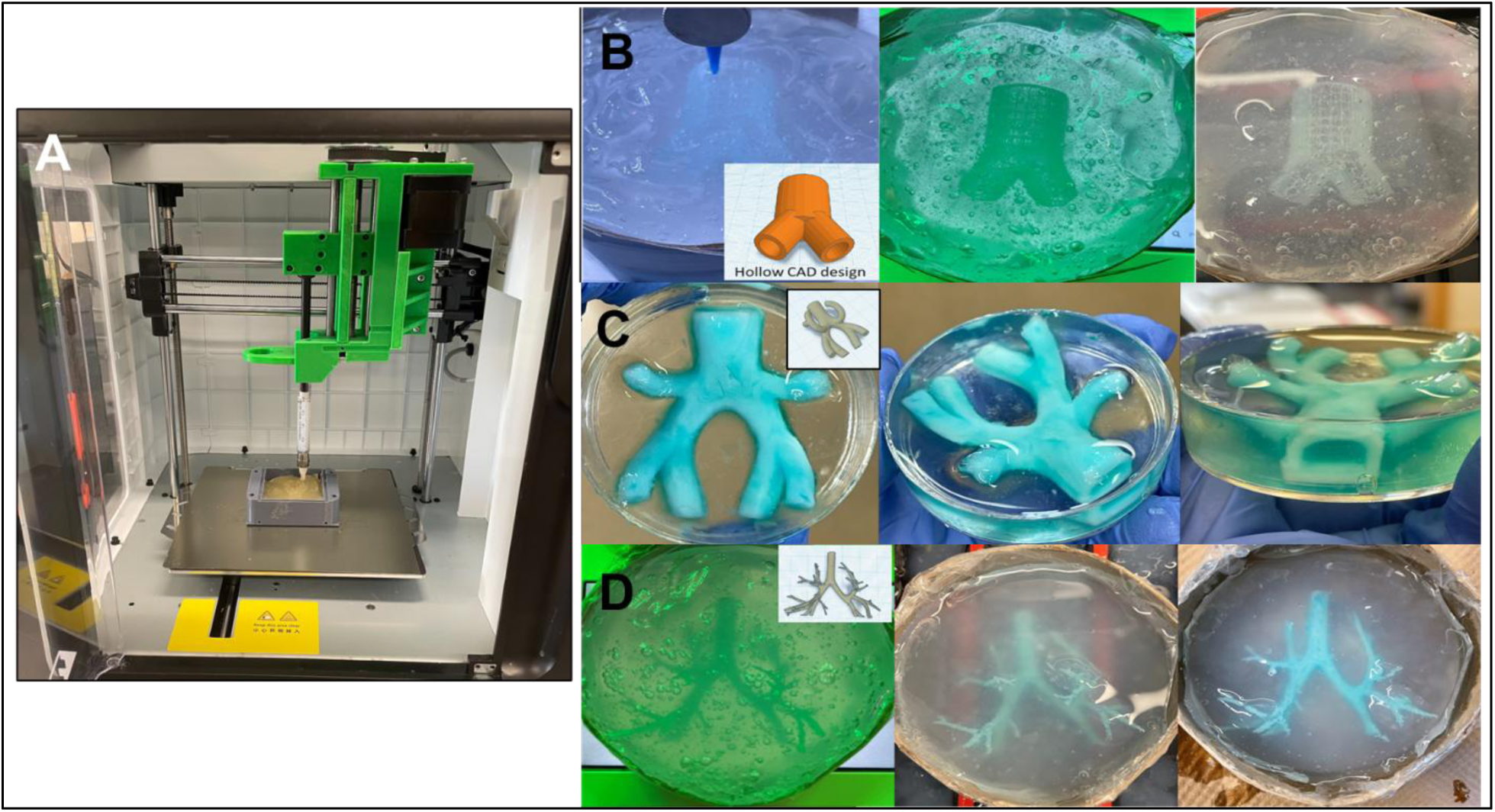
Simplified and CT-derived hollow airway structures 3D bioprinted using AW-dECM + Cellink-RGD bioink and the FRESH approach. **3A:** Image of FlashForge Adventurer 3D printer converted into 3D bioprinter. **3B:** 3D bioprinted airway-like structure printed into support bath. **3C:** 3D bioprinted simplified, hollow airway structure. Blue food dye added for visualization. **3D:** 3D bioprinted airway structure derived from a pediatric CT-scan. Blue food dye added for visualization.

### 2.3. Primary airway epithelial cells exhibit enhanced viability, adhesion and spreading on AW-dECM Cellink-RGD bioinks *in vitro*

To assess *in vitro* biocompatibility, Cellink bioinks conjugated with RGD peptides were combined with 30 mg/mL AW-dECM as described previously. Human primary bronchial epithelial cells successfully adhered to bioinks and exhibited excellent spreading and viability by Day 3 (**Figure 4A**). Additionally, growth and spreading were further enhanced by Day 14 as indicated by Calcein AM/Ethidium bromide staining, demonstrating the excellent *in vitro* biocompatibility of these bioinks with airway epithelial stem cells (**Figure 4A**).

**Figure 4:**
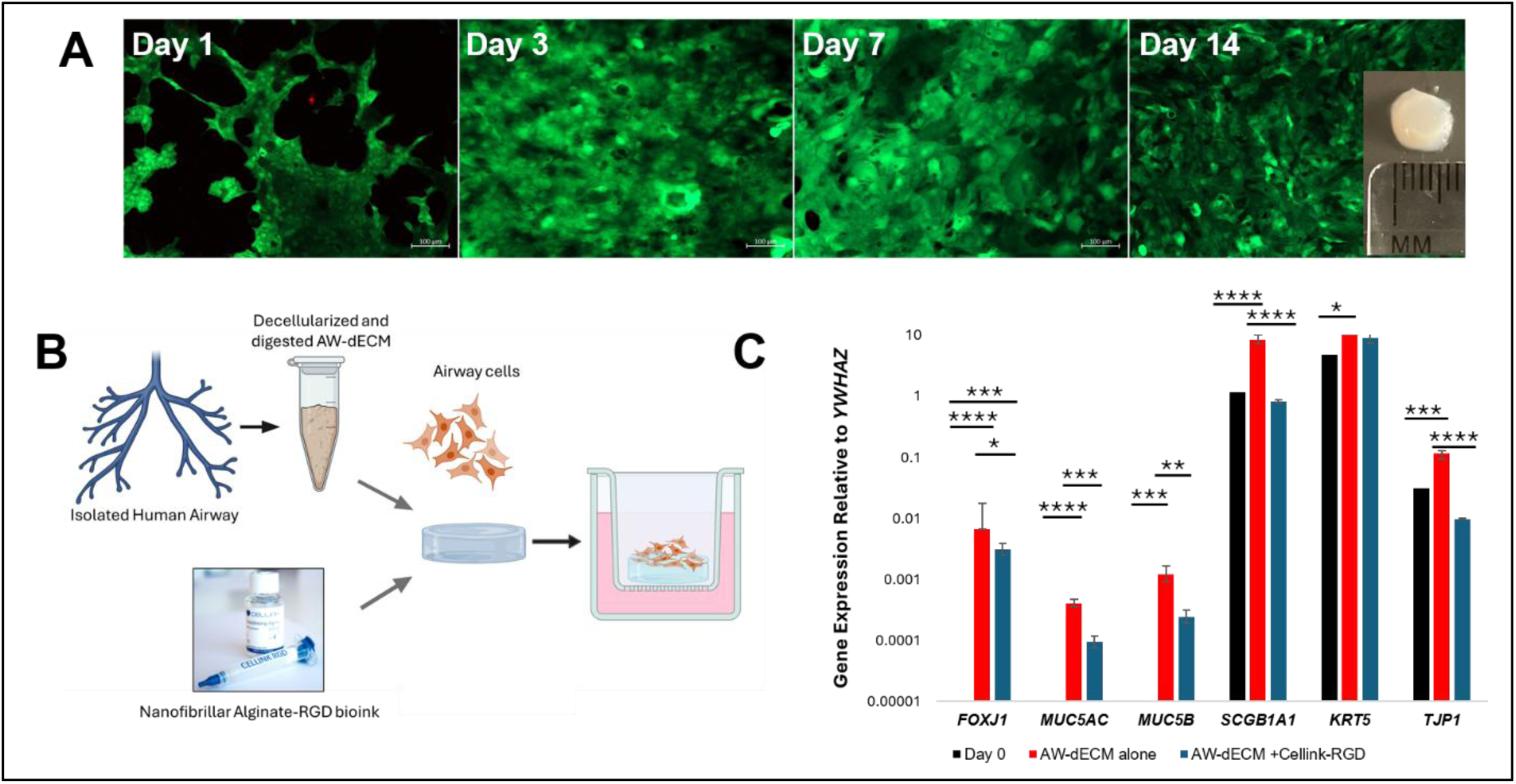
Viability and characterization of primary human airway epithelial cells seeded on AW-dECM + Cellink-RGD bioinks cultured at Air Liquid-Interface over 28 Days. **4A:** Live/dead analysis of primary human airway epithelial cells seeded on top of 30 mg/mL AW-dECM + Cellink-RGD bioink over 14 Days, scale bars= 100 µm. **4B:** Schematic of cell seeding experiment. **4C**: qPCR analysis of *FOXJ1, MUC5AC, MUC5B, SCGB1A1, KRT5* and *TJP1* expression in primary human airway epithelial cells (P2) seeded on 30 mg/mL AW-dECM + Cellink-RGD bioinks. Mean ± SD. p<0.05 *; p<0.01 **; p<0.001 ***; p<0.0001 ****, n=4. Data calculated using delta-delta Ct method. Statistical analysis performed on fold gene expression values (2^-(ΔΔCt).

### 2.4. AW-dECM polymer-blended bioinks conjugated with RGD support the differentiation of seeded human airway epithelial cells to mucociliary and secretory phenotypes over 28 Days

qPCR analysis revealed the successful differentiation of primary human airway cells seeded on bioinks to mucociliary and secretory phenotypes over 28 Days (**Figure 4B + 4C**). Results were compared to Day 0 cells and airway cells seeded on 30 mg/mL AW-dECM alone for 28 days. Results revealed a statistically significant increase in *FOXJ1* expression in primary airway cells seeded on AW-dECM alone and AW-dECM + Cellink-RGD compared to Day 0 (p≤0.0001 and p≤0.001, respectively) (**Figure 4C**). There was also a statistically significant increase in *MUC5AC* and *MUC5B* expression across both conditions compared to Day 0. For *MUC5AC* expression, there was a statistically significant increase in AW-dECM + Cellink-RGD vs Day 0 samples (p≤0.05) and AW-dECM alone vs. Day 0 samples (p≤0.0001) (**Figure 4C**). For *MUC5B* expression, there was a statistically significant increase in AW-dECM + Cellink-RGD vs. Day 0 samples (p≤0.05) and AW-dECM alone vs. Day 0 samples (p≤0.001) (**Figure 4C**). In terms of *SCGB1A1* expression, there was no significant difference in Day 0 vs. AW-dECM + Cellink-RGD samples, however there was a statistically significant increase in samples seeded on AW-dECM alone vs. Day 0 and AW-dECM + Cellink-RGD (p≤0.0001) (**Figure 4C**). There was also a statistically significant increase in basal cell expression (*KRT5*) in cells seeded on AW-dECM alone compared to Day 0 samples (p≤0.05), however no difference in expression was seen between cells at Day 0 and cells seeded on AW-dECM + Cellink-RGD indicating basal cell phenotype was maintained over the 28 day period (**Figure 4C**). Overall, *TJP1* expression was elevated on AW-dECM alone compared to Day 0 and AW-dECM + Cellink-RGD samples.

### 2.5. Acellular AW-dECM Cellink-RGD bioinks implanted subcutaneously in a rat model over one month demonstrate limited degradation and are biocompatible

Sprague-Dawley rats implanted with bioprinted Cellink-RGD and AW-dECM Cellink-RGD bioink samples over 30 days demonstrated excellent health with no signs of swelling or surgical site infection. Following tissue harvest on Days 3, 14 and 30, macroscopic evaluation of harvested implantation sites revealed no signs of necrosis, edema, infection or tissue rejection (**Supplemental Figure 3A-3G**). On Day 3, Cellink-RGD bioink samples were largely mobile within the skin, indicating a lack of encapsulation within the subcutaneous wall and muscle layer. AW-dECM Cellink-RGD bioinks were immobile on Day 3 and gross macroscopic evaluation of harvested implantation sites revealed the presence of a pocket-shaped hole within the subcutaneous wall where the bioinks were deposited (**Supplemental Figure 3G**). By Days 14 and 30, both types of bioinks were encapsulated within the tissue as indicated by a lack of mobility within the skin. At the microscopic level, H & E staining of both bioink types at all time points revealed a lack of tissue integration (**Figure 5**), however, cellular infiltration was evident at the bioink-tissue interface (**Supplemental Figure 4**). Measurement of this layer at Days 14 and 30 in Cellink-RGD and AW-dECM Cellink-RGD bioinks revealed no significant differences in thickness between both samples and at both time points (**Supplemental Figure 4B**).

**Figure 5:**
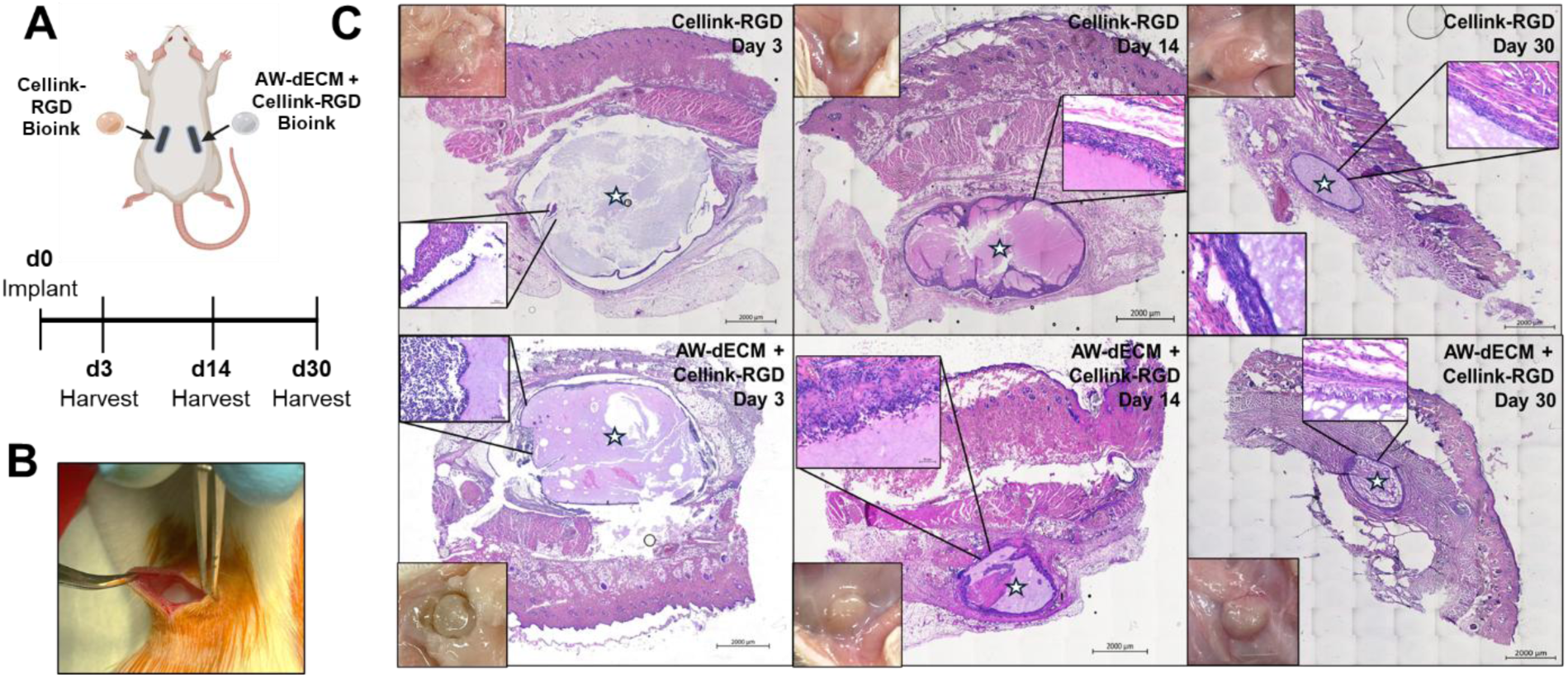
Representative H & E Staining of Acellular Cellink-RGD and AW-dECM +Cellink-RGD bioinks implanted subcutaneously in immune competent Sprague Dawley rats over 30 Days. **5A:** Outline of experimental approach and timeline, n=8 rats/time point. **5B**: Imaging depicting placement of bioink into subcutaneous tissue pocket. **5C**: Representative H & E staining of implanted bioinks at different time points. White star symbol denotes subcutaneously implanted bioink. Inset H & E images of bioink edge = 50 µm magnification. Inset macroscopic images in upper left (**top panels**) and lower left (**bottom panels**) depict bioink following tissue harvest at **Day 3**, **Day 14** and **Day 30**.

Masson’s Trichrome staining demonstrated the presence of a typical foreign body response evident by the formation of a fibrotic capsule surrounding the implanted bioink (**Figure 6**). Immunofluorescent staining of Day 3 bioink samples with a neutrophil-specific marker Myeloperoxidase, confirmed the presence of acute inflammatory cells on the bioink surface and in the surrounding tissues following bioink implantation (**Supplemental Figure 5**), that resolved by Day 14 (**Supplemental Figure 6**). Additionally, immunofluorescent staining of the cells in proximity to the bioink revealed the presence of M1 and M2 macrophages at Days 14 and 30 (**Figure 7**). Comparison of the storage and loss moduli of Day 0 vs. Day 30 bioinks demonstrated no significant difference in values between both Cellink-RGD alone and AW-dECM + Cellink-RGD, suggesting minimal degradation occurred over the 30-day period following transplantation (**Figure 8A + 8B**). To further support this assessment, measurement of Day 30 bioinks compared to Day 0 revealed no change in diameter over time for both types of bioinks (**Figure 8C + 8D**).

**Figure 6:**
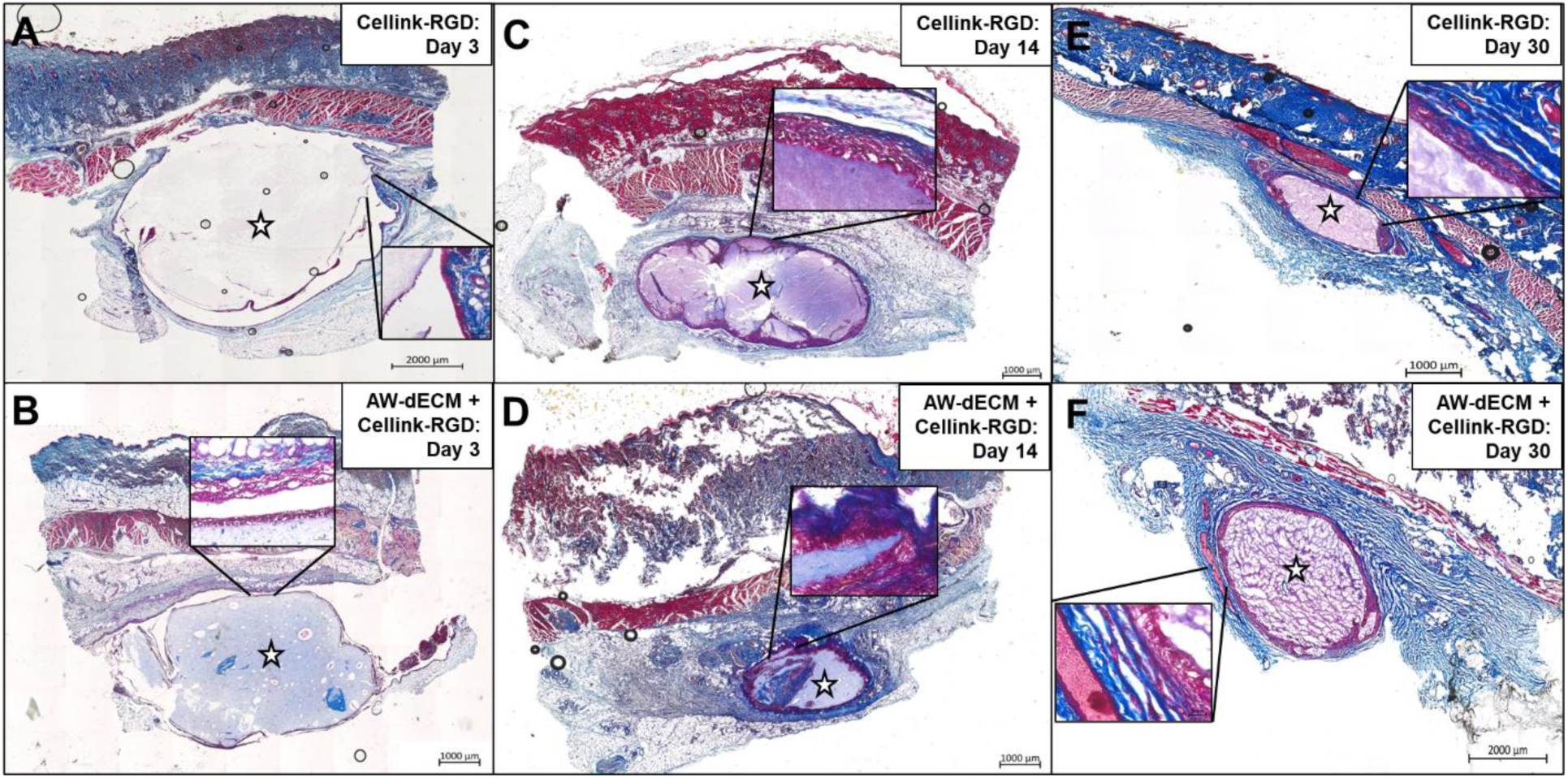
Representative Masson’s Trichrome staining of acellular Cellink-RGD and AW-dECM +Cellink-RGD bioinks implanted subcutaneously in immune competent Sprague Dawley rats over 30 days. **6A + 6B:** Cellink-RGD and AW-dECM + Cellink-RGD on Day 3. **6C + 6D:** Cellink-RGD and AW-dECM + Cellink-RGD on Day 14. **6E + 6F**: Cellink-RGD and AW-dECM + Cellink-RGD on Day 30. Star symbol denotes subcutaneously implanted bioink. Inset images, scale bar= 50 µm.

**Figure 7:**
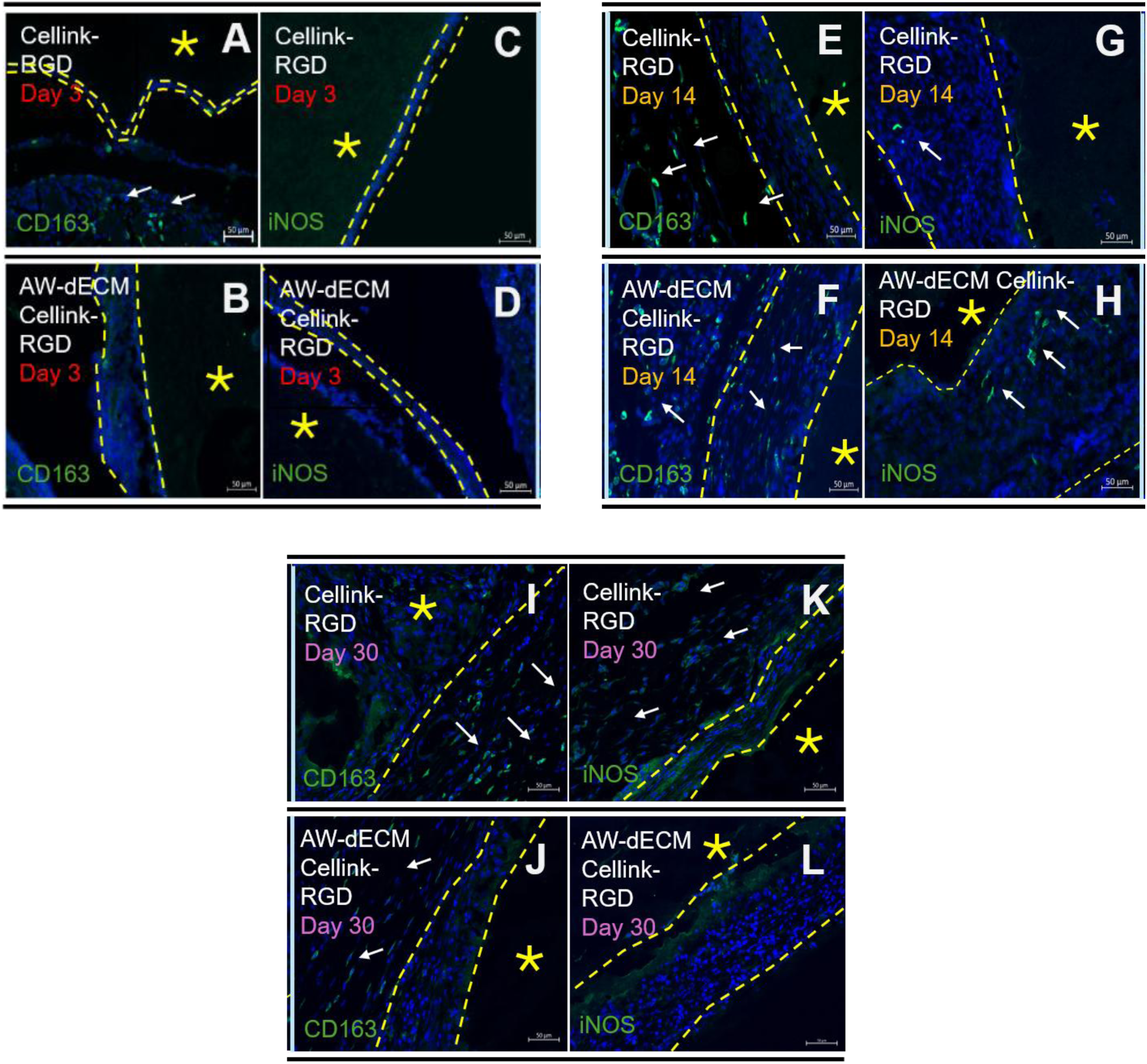
Immunofluorescent analysis of macrophage populations at bioink-tissue interface at Days 3, 14 and 30. **7A + 7B:** CD163+ staining of cells at bioink-tissue interface in Cellink-RGD and AW-dECM Cellink-RGD samples on Day 3. **7C + 7D:** iNOS+ staining of cells at bioink-tissue interface in Cellink-RGD and AW-dECM Cellink-RGD samples on Day 3. **7E + 7F:** CD163+ staining of cells at bioink-tissue interface in Cellink-RGD and AW-dECM Cellink-RGD samples on Day 14. **7G + 7H:** iNOS+ staining of cells at bioink-tissue interface in Cellink-RGD and AW-dECM Cellink-RGD samples on Day 14. **7I + 7J:** CD163+ staining of cells at bioink-tissue interface in Cellink-RGD and AW-dECM Cellink-RGD samples on Day 30. **7K + 7L:** iNOS+ staining of cells at bioink-tissue interface in Cellink-RGD and AW-dECM Cellink-RGD samples on Day 30. Yellow dotted lines demarcate bioink-tissue interface, yellow star denotes bioink sample. White arrows indicate cells with positive staining. Scale bars= 50 µm.

**Figure 8:**
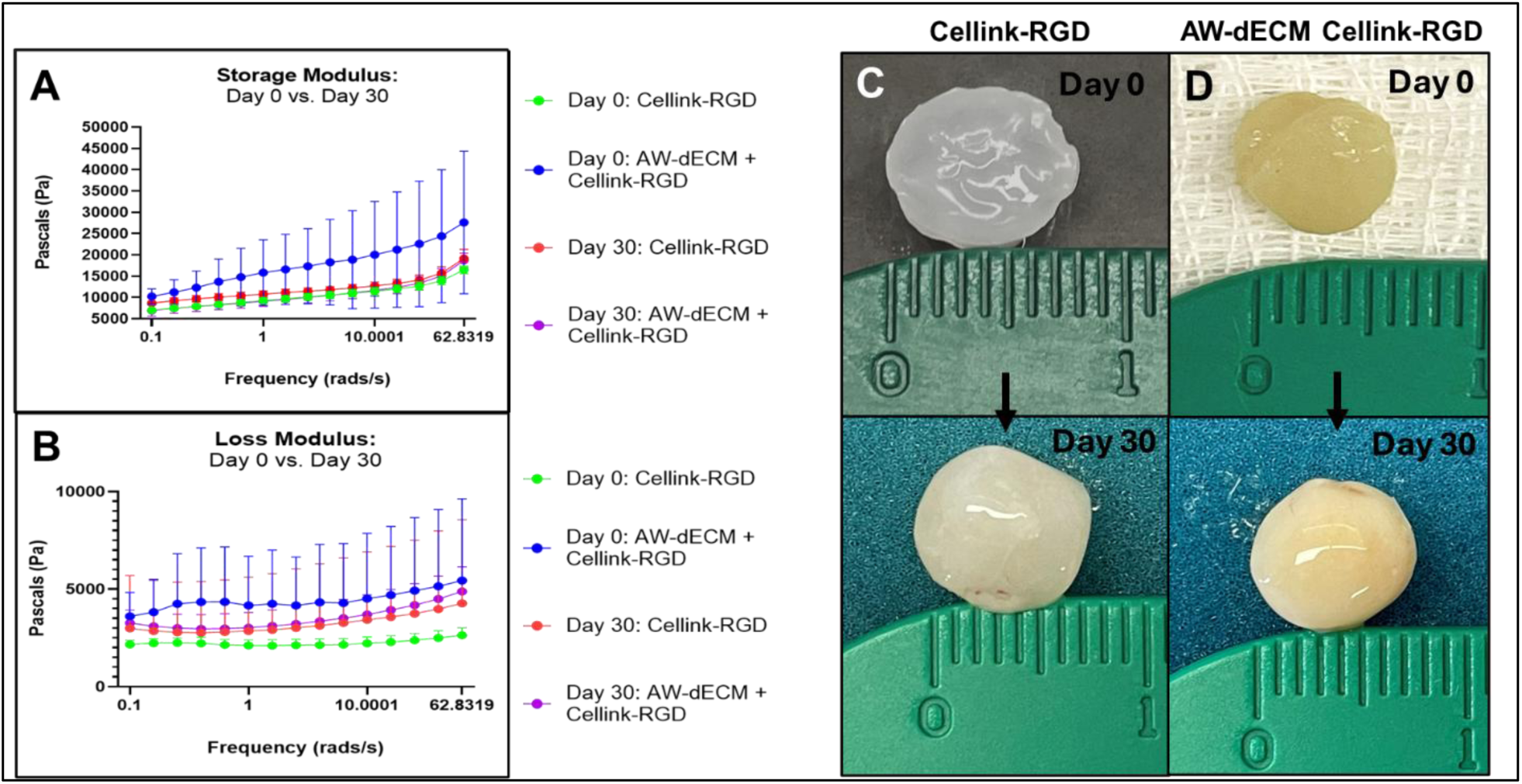
Mechanical assessment and degradation of Cellink-RGD and AW-dECM Cellink-RGD bioinks following subcutaneous implantation in rats over 30 Days. **8A:** Storage Modulus in Pascals (Frequency Sweep) of Cellink-RGD vs. AW-dECM Cellink-RGD bioinks at Day 0 vs. Day 30 of SQ implantation. **8B**: Loss Modulus in Pascals (Frequency Sweep) of Cellink-RGD vs. AW-dECM Cellink-RGD bioinks at Day 0 vs. Day 30 of SQ implantation. Data represented as mean ± SD, n=6 bioinks/time point (Day 0), n=4 bioinks/time point (Day 30). **8C + 8D**: Measurement of Cellink-RGD and AW-dECM Cellink-RGD bioinks at Day 0 vs. Day 30.

## DISCUSSION

Despite significant advances in airway engineering, existing bioinks are limited in their ability to simultaneously reproduce tissue-specific biochemical cues, native-like mechanics, and hollow architecture required to generate physiologically relevant proximal airway constructs. Here, we address this challenge through the development of a human airway-derived decellularized extracellular matrix (AW-dECM)-based polymer blended bioink that combines tissue-specific biological signaling with polymer-enhanced printability and structural stability. This approach enables the fabrication of simple, hollow cylinders as well as more complex airway structures derived from a CT scan. Primary human airway epithelial cells seeded on top of bioinks and grown at air-liquid interface over 28 days exhibited excellent biocompatibility, viability, adhesion and spreading *in vitro*. Cells also successfully differentiated to mucociliary and secretory phenotypes over 28 days. *In vivo* studies using immunocompetent rats demonstrated a lack of necrosis and infection at the surgical site following acellular bioink implantation over 30 days. Collectively, these findings establish a strategy for recreating the airway-specific microenvironment while also providing a foundation for more translational *ex vivo* airway models and regenerative airway therapies.

Compared to AW-dECM alone, AW-dECM blended with Cellink-RGD exhibited enhanced mechanical properties. The storage moduli (∼8 kPa) of the polymer blended bioinks more closely reflects values reported for native lung tissue (∼5-10 kPa) compared to AW-dECM alone (∼100 Pa), enabling the successful bioprinting of hollow airway-like structures with improved shape fidelity ^44^. For many ECM-based tissue sources, the decellularization and hydrogel/bioink formation process renders the tissue less mechanically stable due to the breakdown of collagen telopeptides during the pepsin digestion step ^45^. This makes the materials unsuitable for extrusion-based bioprinting as the resultant biomaterial is much softer and subsequently unable to hold its shape following deposition into a support bath or culture dish. The addition of natural or synthetic polymers enhances the mechanical properties of the bioinks significantly, allowing for ease of bioprinting and increased shape fidelity. In this study, commercially available alginate conjugated to RGD was chosen as an optimal polymer as it is derived from natural resources, is inexpensive, and demonstrates enhanced biocompatibility and cell adhesion compared to alginate alone and other synthetic bioinks such as PEG and PLGA ^46^.

Based on our *in vitro* findings, human airway epithelial cells can successfully differentiate to mucociliary and secretory phenotypes when seeded on AW-dECM Cellink-RGD bioinks, albeit at lower levels compared to AW-dECM alone. This may be due to the inherent reduced degradability of alginate compared to native ECM, which could have prevented the cells from fully breaking down and remodeling the underlying matrix ^47^. This limitation could be addressed through incorporation of MMP-sensitive peptides within the bioinks, to encourage tissue breakdown and appropriate regeneration ^48^. In one study, MMP2-sensitive peptides incorporated into alginate hydrogels laden with human mesenchymal stem cells promoted increased degradation and tissue integration following subcutaneous implantation in xenograft mouse models ^48^.

Our *in vivo* results of subcutaneously implanted bioinks demonstrated a lack of necrosis and infection in immunocompetent Sprague Dawley rats over 30 days. After 30 days, the bioinks maintained their original size and mechanical properties as indicated by macroscopic and rheological evaluation of harvested bioinks at Day 0 vs. Day 30. However, a typical foreign body response (FBR) to both Cellink-RGD alone and AW-dECM Cellink-RGD bioinks was evident by Days 14 and 30. Histological staining revealed the development of a well-formed fibrous capsule in both conditions by the end of one month. Limited data currently exists related to the immunogenicity of Alginate-RGD biomaterials *in vivo* as most studies primarily focus on its *in vitro* biocompatibility ^49, 50^. Therefore, this study provides pertinent information related to the host immune response of commercially available alginate-RGD biomaterials and indicates that its composition requires further modification to ensure appropriate tissue integration and biocompatibility prior to use in clinical applications.

Another study also investigated the utility of alginate-blended (no RGD) ***bulk lung*** bioinks in 3D bioprinting lung-specific tissue. Researchers found that the material exhibited excellent printability and biocompatibility of embedded and seeded primary human epithelial cells, however the samples were subcutaneously implanted in a mouse model of *transplant immunosuppression* ^51^. The preferred method is to assess biocompatibility in immunologically competent subjects, as the use of immunosuppressive medications following transplantation is not ideal and can lead to a heightened risk of infections, malignancy and organ toxicity ^52^. Therefore, the current study informs on the host response to Cellink RGD alone and AW-dECM + Cellink RGD in immunocompetent hosts. Although these materials are rated as immunologically inert and biocompatible, they still elicit a typical *in vivo* foreign body response, and further modifications will be required to enhance their clinical applicability and safety.

Pre-clinical studies have suggested that incorporating biomaterials with anti-inflammatory bioactive peptides or cytokines such as IL-4 that promote macrophage polarization to M2 phenotypes, hold promise in mitigating the FBR ^53–55^. Our assessment of the cell types within the fibrous capsule at Day 30 indicated the presence of both M2 and M1 macrophages. This aligns with current data that demonstrates the involvement of macrophages in regulating the FBR to implanted hydrogels ^56^. Future studies will focus on incorporating these anti-inflammatory components within AW-dECM Cellink-RGD bioinks to promote improved biocompatibility and integration. Additionally, studies suggest that bioink stiffness can also influence the subsequent immune response following implantation. One study demonstrated that stiffer hydrogels (10 kPa) promoted a more pro-inflammatory macrophage phenotype compared to softer hydrogels (<2 kPa) ^57^. Therefore, the stiffness of this bioink formulation could be modulated by decreasing the crosslinking time to create a slightly softer substrate. Limitations of this study include the use of xenogeneic materials to assess host immune response and overall biocompatibility. Moving forward, decellularized *rat* airway tissue combined with Cellink-RGD will be tested in immunocompetent animals to determine if the host response and tissue integration is improved. Additionally, the host response following orthotopic implantation will need to be assessed as the subcutaneous space differs drastically in terms of immune response to foreign materials.

In conclusion, this study demonstrates the potential of human AW-dECM + Cellink-RGD bioinks for 3D bioprinting simple and complex hollow airway structures with strong mechanical and biochemical properties, while also providing insight into host immune responses to alginate - RGD biomaterials alone and combined with human-derived decellularized airway tissue. This information will be pertinent in the improved design of biocompatible materials useful for *ex vivo* airway modeling and tissue engineering applications.

**Supplemental Table 1:**
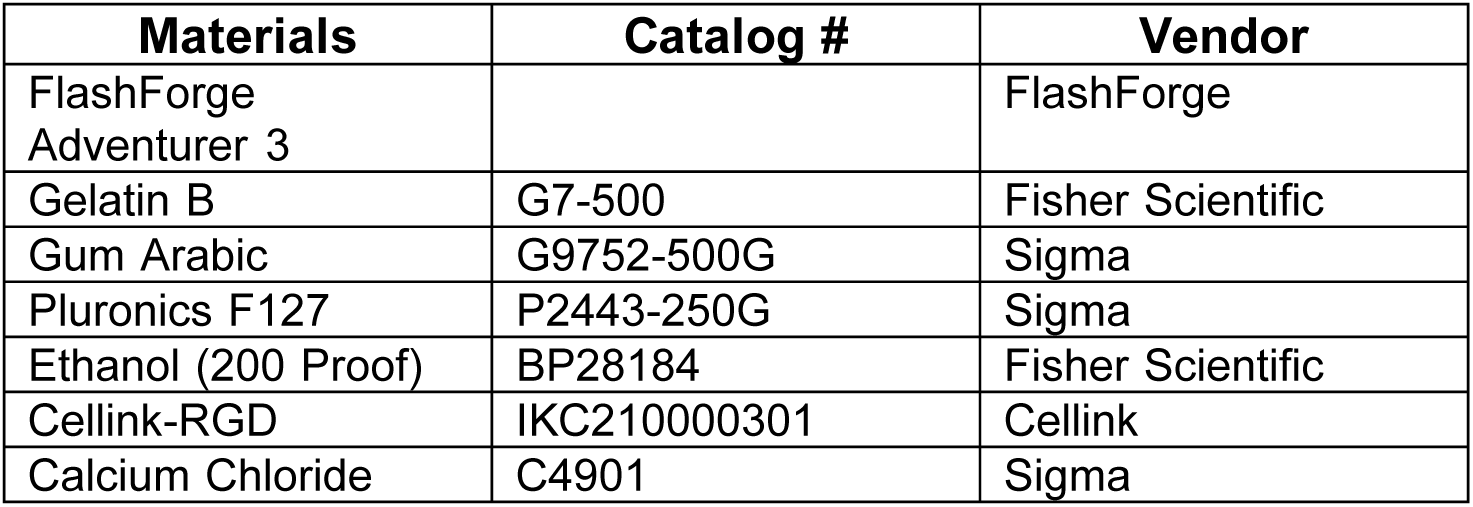
Materials used for 3D Bioprinting.

**Supplemental Table 2:** List of Biorad Primers used for RT-qPCR.

| Primer Name | Unique Assay ID |
| --- | --- |
| *YWHAZ* | qHsaCID0013897 |
| *B2M* | qHsaCID0015347 |
| *FOXJ1* | qHsaCID0016777 |
| *MUC5B* | qHsaCID0011690 |
| *MUC5AC* | qHsaCID0017663 |
| *SCGB1A1* | qHsaCID0018013 |
| *TP63* | qHsaCID0036332 |
| *KI67* | qHsaCID0011882 |
| *RSPH4A* | qHsaCID0017043 |
| *KRT5* | qHsaCED0047798 |

**Supplemental Table 3:** List of Primary and Secondary Antibodies used for IF Analysis.

| Antibodies | Vendor |
| --- | --- |
| 1°: CD163 Rabbit | Thermofisher |
| 1°: iNOS, Rabbit | Thermofisher |
| 1°: Myeloperoxidase, Rabbit | Thermofisher |
| 2°: Goat Anti-Rabbit | Thermofisher |

## CRediT authorship contribution statement

**Heather Wanczyk**: Writing – review & editing, Writing – original draft, Methodology, Investigation, Formal analysis, Data curation, Conceptualization. **Nina Kosciuszek**: Surgical assistance, Writing-Review and editing. **Joanne Walker**: Writing – review & editing, Investigation, Data curation. **Daniel Weiss**: Writing – review & editing, Supervision, Project administration, Methodology, Investigation, Funding acquisition, Conceptualization. **Christine Finck**: Writing – review & editing, Supervision, Project administration, Methodology, Investigation, Funding acquisition, Conceptualization.

## Acknowledgements

We would like to acknowledge the generous donation of human primary airway cells from the Ryan Lab at the University of Iowa that were used in this study.

## Statements and declarations

### Ethical considerations

Ethical approval was not required.

### Consent to participate

Not applicable.

### Consent for publication

Not applicable.

### Data availability

Data will be made available on request.

