## Supplemental Figure for "Recreating the Native Airway Microenvironment Using Tissue-Specific Extracellular Matrix Bioinks for Proximal Airway Engineering"

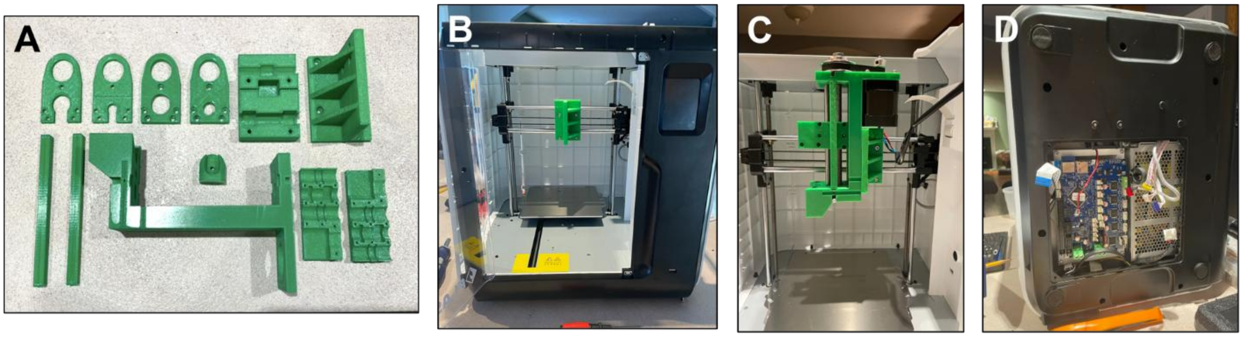


**Supplemental Figure 1**: **Fabrication of 3D bioprinter from FlashForge FDM 3D desktop printer. S1A**: 3D printed PLA parts for fabricating bioprinter. **S1B**: Flashforge printer with original hot-end nozzle removed. **S1C**: Desktop printer with 3D printed Replistruder syringe parts assembled. **S1D**: Image of Duet wifi board replacement.

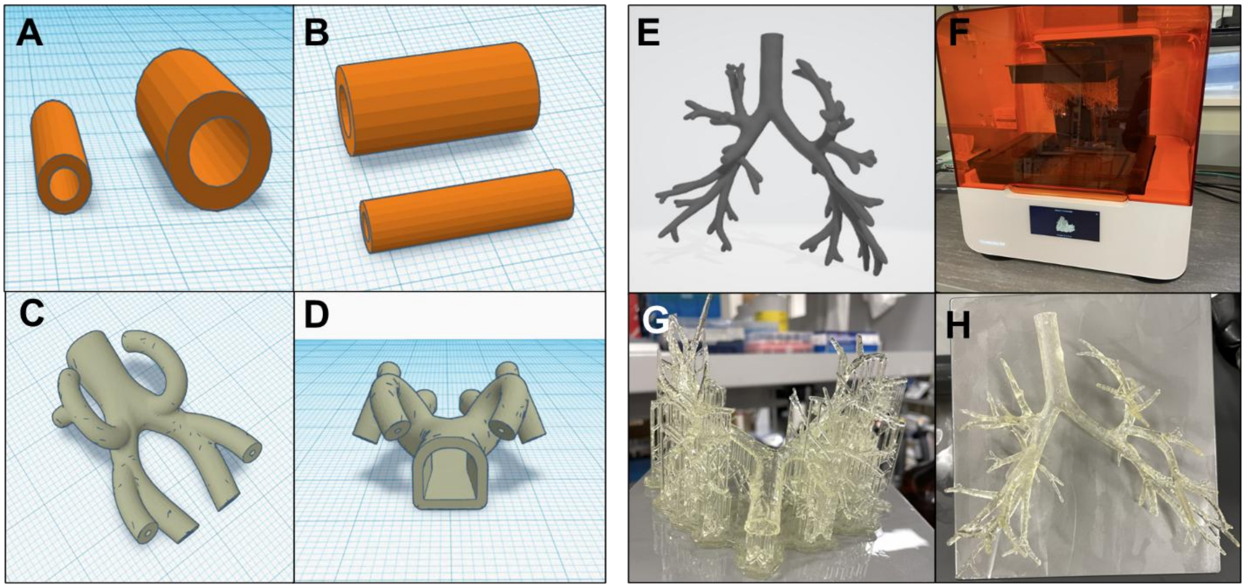


**Supplemental Figure 2**: **CAD files used for 3D bioprinting hollow cylinders and simplified or complex hollow airway structures. S2A+S2B**: Hollow cylinders (5-10 mm diameters, 1 mm or 2 mm thickness) mimicking size of pediatric trachea. **S2C + 2D**: Simplified, branched hollow airway structure. **S2E**: Pediatric airway structure derived from CT-scan. **S2F**: Stereolithography (SLA) 3D printer used for testing printability of hollow pediatric airway structure derived from CT-scan. **S2G + S2H**: Hollow pediatric airway 3D printed using resin (with supports and without).

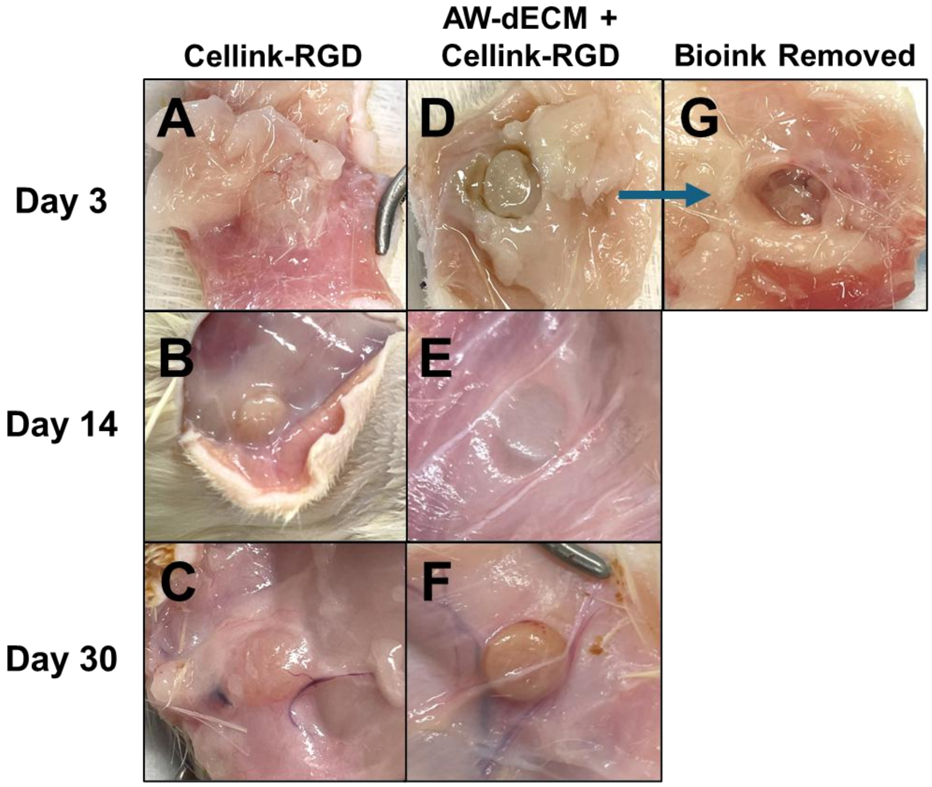


**Supplemental Figure 3: Representative macroscopic images of harvested bioink samples at days 3, 14 and 30. S3A, S3B, S3C:** Cellink-RGD samples at three different time points. **S3D, S3E, S3F:** AW-dECM + Cellink-RGD samples at three different time points**. S3G:** Pocket formed in tissue by AW-dECM + Cellink-RGD bioink at Day 3.

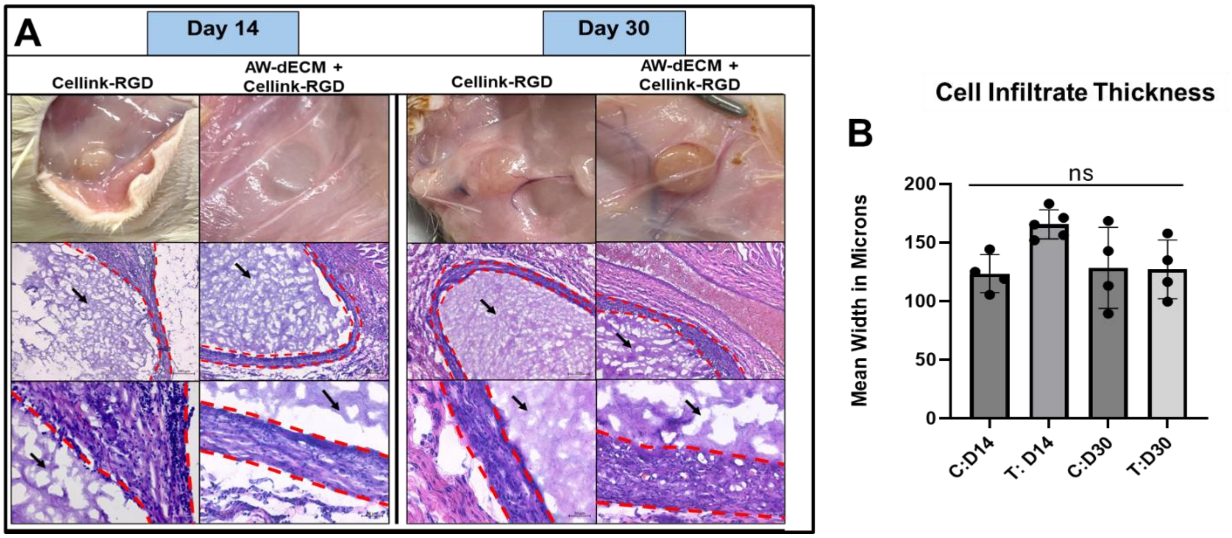


**Supplemental Figure 4: Representative H&E images of bioink-tissue interface. S4A:** H&E images of bioink-tissue interface at Day 14 and Day 30 revealing formation of fibrotic capsule in Cellink-RGD and AW-dECM + Cellink-RGD bioink samples. Black arrow denotes bioink. Red dotted line delineates cellular infiltrate at bioink-tissue interface. **S4B:** Measurement of cellular infiltration/fibrotic capsule thickness at bioink-tissue interface in microns. n=4 biological samples, 6 images/sample, 10 measurements/image. Data represented as mean ± SD. Statistical significance measured using One-way Anova, Tukey’s Multiple Comparisons.

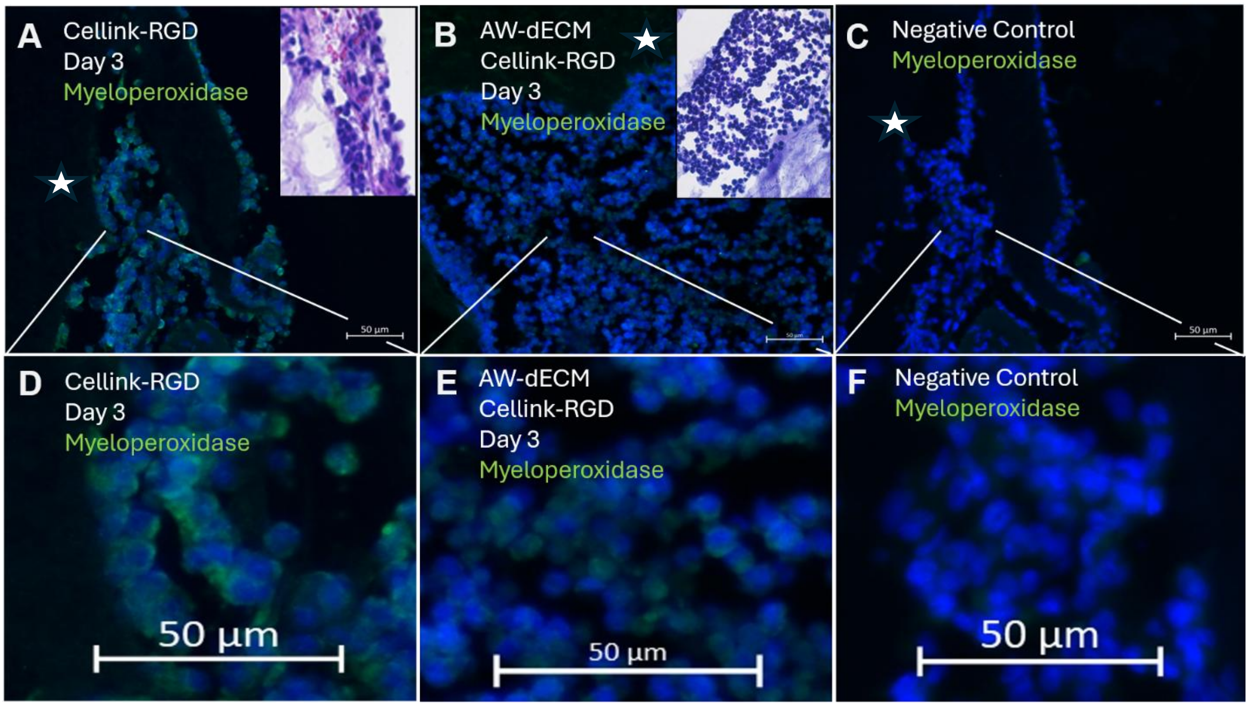


**Supplemental Figure 5: Immunofluorescence analysis of immune cells at Bioink-tissue interface on Day 3. S5A:** Representative myeloperoxidase staining of cells surrounding AW-dECM +Cellink-RGD bioink samples at Day 3. **S5B:** Representative myeloperoxidase staining of cells surrounding AW-dECM Cellink-RGD bioink sample at Day 3. **S5C:** Negative control. **S5D-S5F:** Corresponding magnified IF images of **S5A-S5C**. White arrows indicate location of bioink samples in proximity to immune cells. Scales bars= 50 µm.

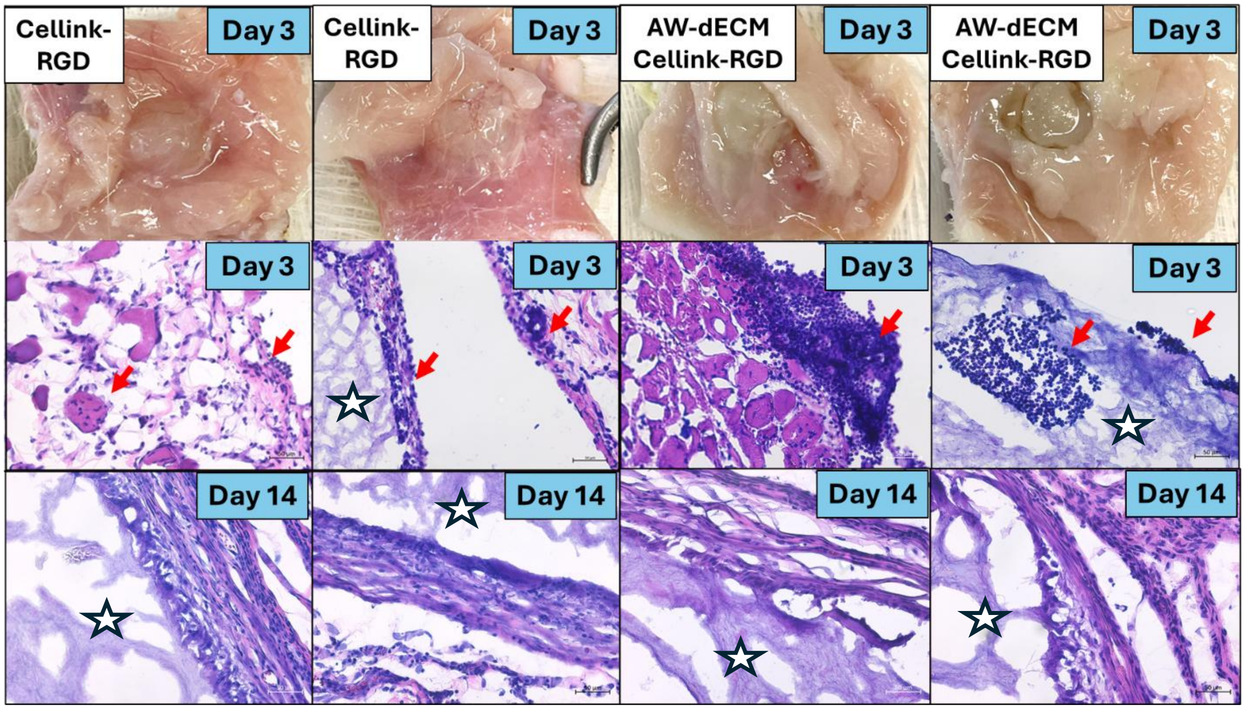


**Supplemental Figure 6: Representative H & E images showing resolution of acute inflammatory response by Day 14 of bioink implantation.** White star indicates location of bioink within tissue. Red arrows indicate inflammatory cells at bioink surface and in surrounding tissue. Scale bars= 50 µm.
